## Supplementary figures and tables for "New insights into the architecture and dynamics of archaella"

### Supplementary material

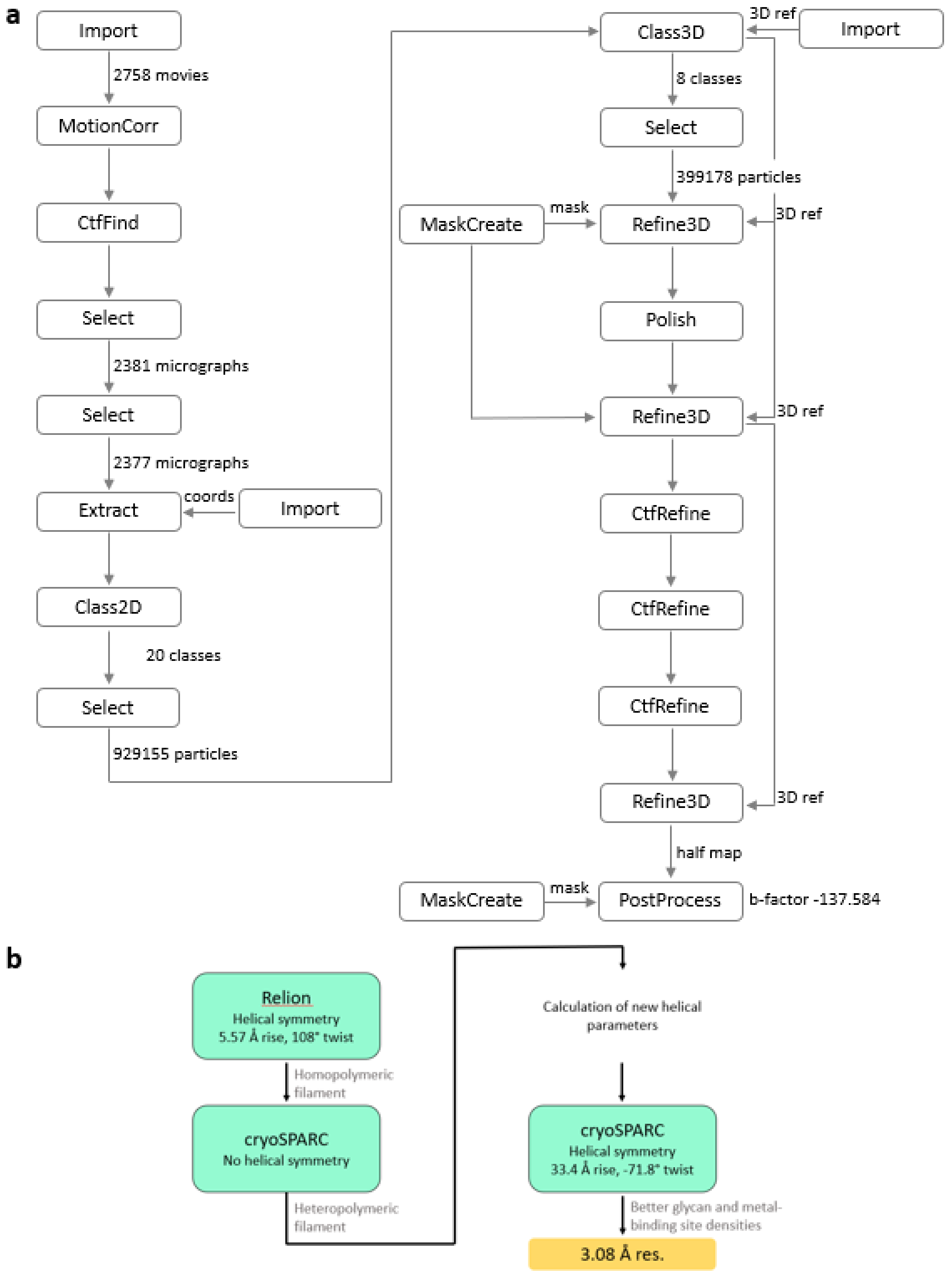


**Suppl. fig. 1|** Data processing flowcharts using Relion 3.1 (a) and cryoSPARC (b).

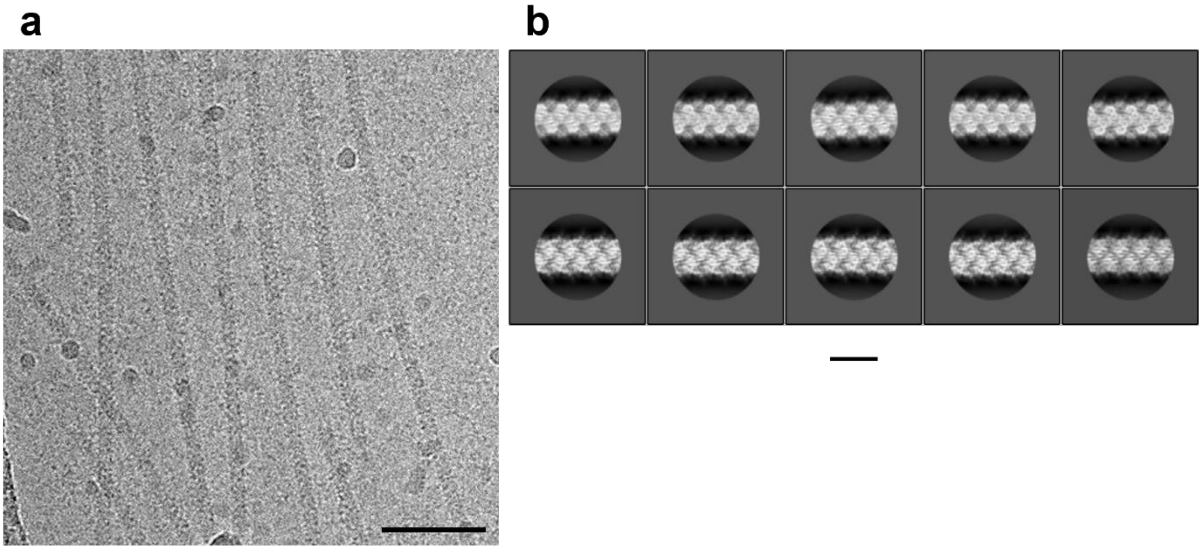


**Suppl. fig. 2| a**, representative cryoEM micrograph of *M. villosus* archaella. **b**, 2D classification examples of polished particles in Relion 3.1. Scale bar in (a), 50 nm; in (b), 10 nm.

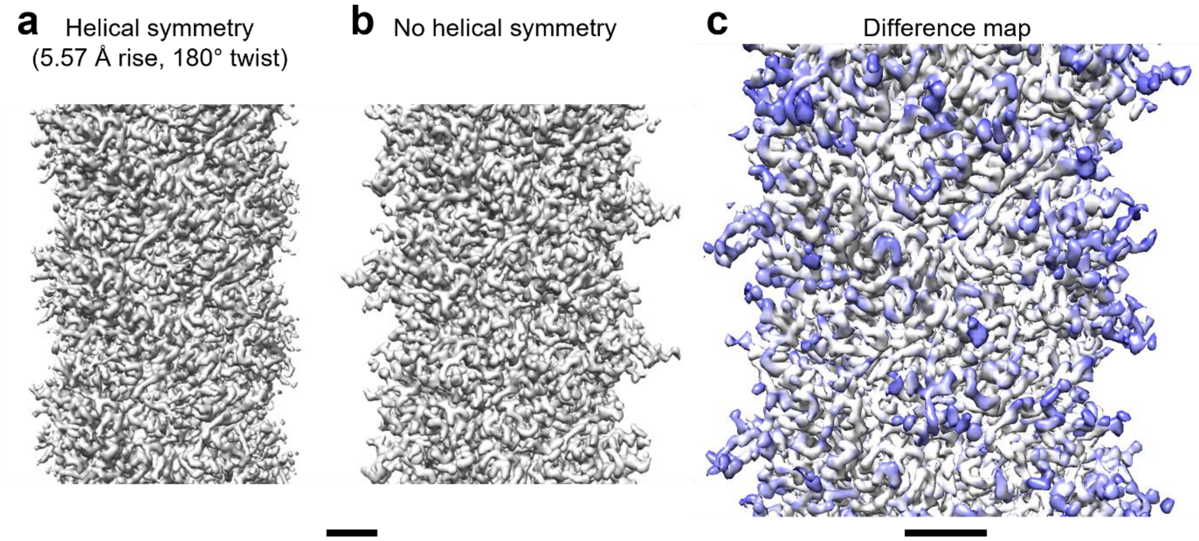


**Suppl. fig. 3|** CryoEM maps of the *M. villosus* archaellum filament obtained applying helical symmetry (5.57 Å rise , 108° twist) (a), without helical symmetry (b) and difference map (c) between the maps shown in (a) and (b). The white areas of the map in (c) are those where the refined maps obtained with and without imposing helical symmetry agree. The blue areas are those better resolved without imposing helical symmetry and that were missing or fragmented in the map with helical symmetry. The areas that benefit from relaxing the helical symmetry (blue) are those occupied by glycans in ArlB1 and 2 and by the “glycosylation loop” in ArlB2. Scale bar 20 Å.

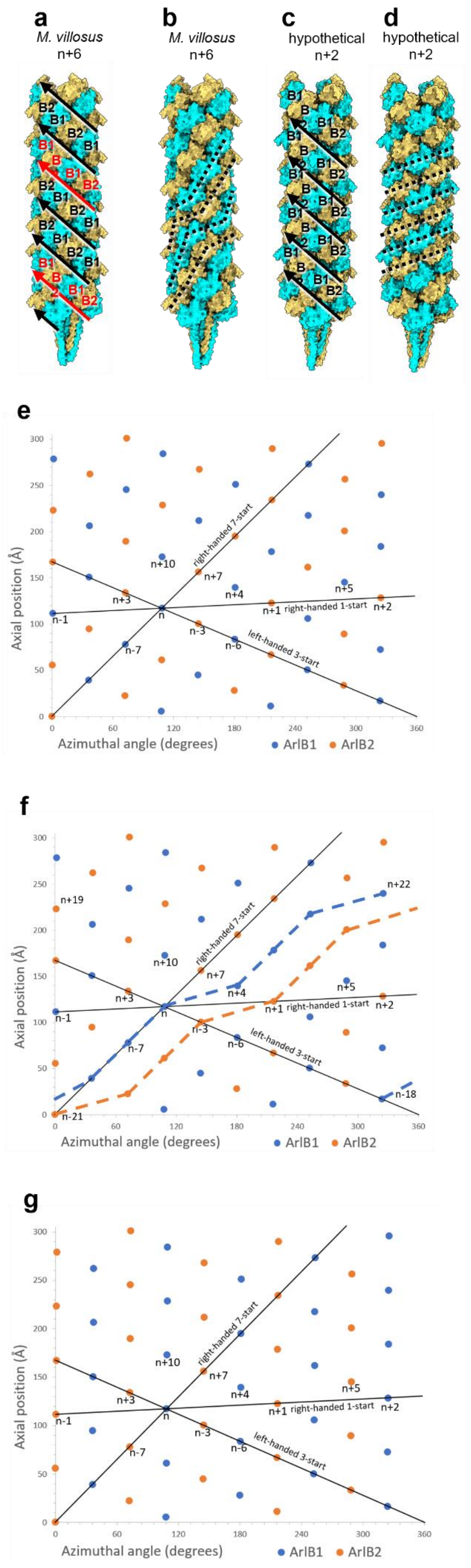


**Suppl. fig. 4| a-d**, the heteropolymeric archaellum from *M. villosus* with n+6 symmetry (a, b) compared with a hypothetical filament with n+2 symmetry (c, d). **a**, in the *M. villosus* filament every third 3-start strand (red) is out of register with respect to the other 3-start helices (black arrows). **b**, subunits of the same type (ArlB1 or ArlB2 only) follow right-handed pseudo strands with broken symmetry. **c**, in a hypothetical n+2 filament, all 3-start strands are perfectly in register. The filament is isotropic and consists of either perfectly alternating or homopolymeric component strands. **d**, the two component subunits form true homopolymeric right-handed 4-start strands.

**e, f**, helical net representations of the arrangement of protein monomers in the *M. villosus* heteropolymeric archaellum, as shown in (a) and (b). The view is from the outside onto the unrolled surface. Similar to homopolymeric archaella, the right-handed one-start helix with helical parameters (108°, 5.6 Å) passes through each protein monomer, the latter are labelled along this helix (e.g., n−1, n, n+1, etc.). The head domains make significant contacts along the right-handed 7-start helical strands (n+7) and the left-handed 3-start strands (n+3), whereas the tail domains also make contacts along the 1-start strand and a 4-start strand. The protein monomers of each type are shown as blue and orange dots, respectively. **e**, the diagram indicates a helical symmetry n+6. The ArlB1-2 subunits alternate along the left-handed 3-start strand, however, they have more complex order along the 1-start or 7-start strands. **f**, nearest monomers of the same type are shown as dashed lines (ArlB1 blue, ArlB2 orange). Eight molecules (including the first and the last near equivalent ones) are related by either n+7 or n+4 contacts and represent a full homopolymeric turn around the filament axis. **g**, a hypothetical heteropolymeric arrangement without screw axis asymmetry, as shown in (c) and (d). Here, the 3-start helix is built up from alternating monomers of each type. It has n+2 helical symmetry, manifested by each 4-start and 10-start strand formed by the monomers of the same type, while monomers of different types alternate along 1-start and 7-start helices.

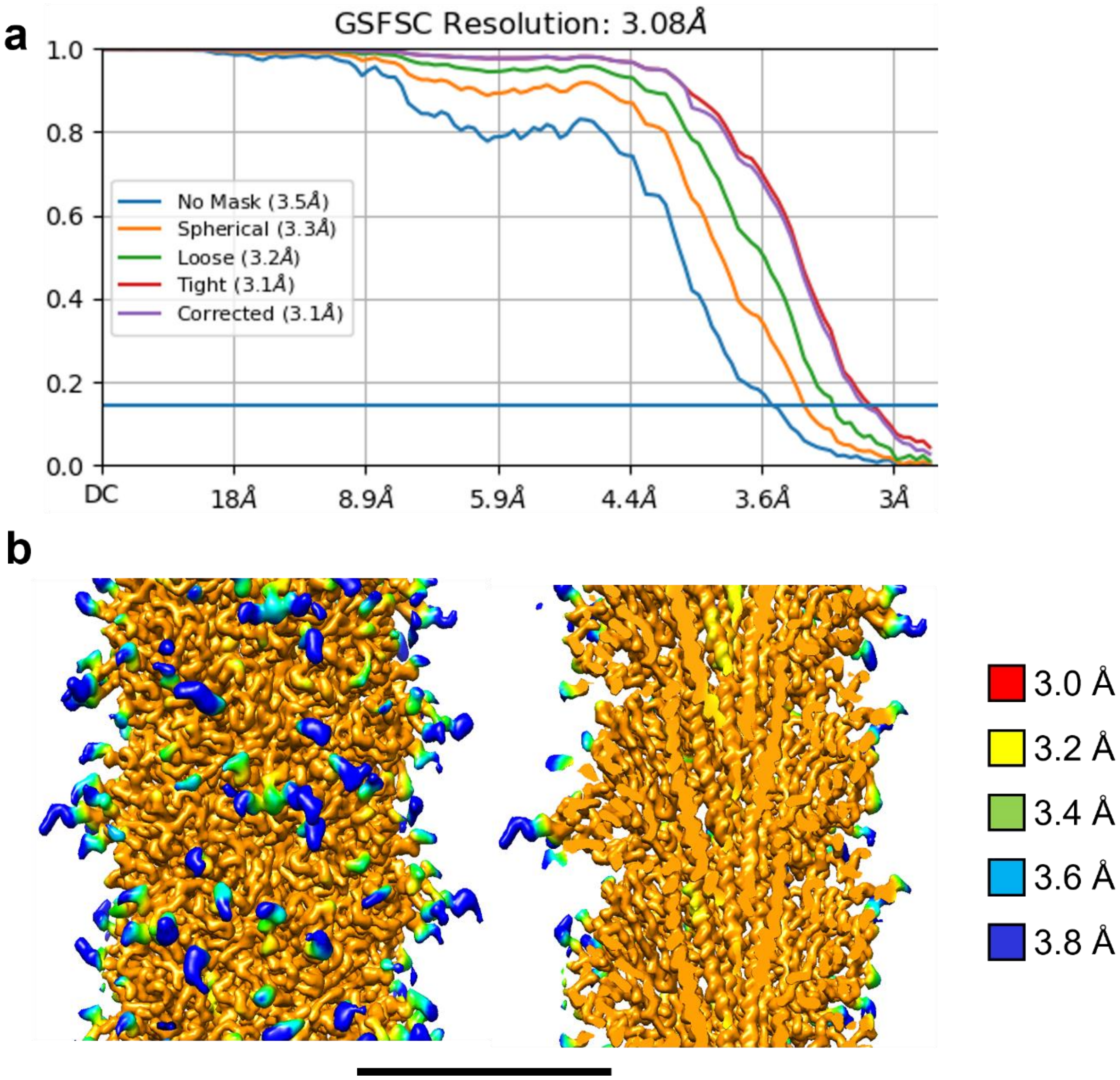


**Suppl. fig. 5| a**, gold-standard FSC and **b**, local resolution estimations for the archaellum map obtained from cryoSPARC 3.0.1 after helical refinement. Scale bar, 50 Å.

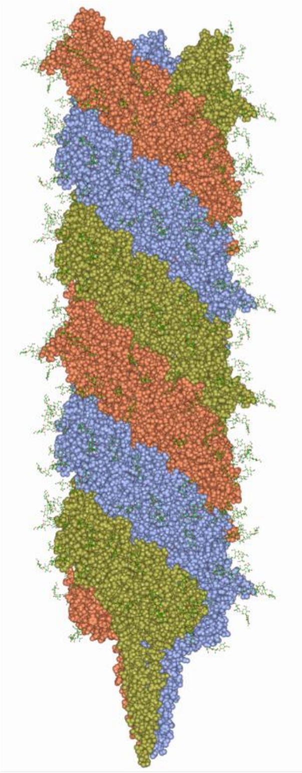

**Suppl. fig. 6|** Atomic model of the *M. villosus* archaellum showing the 3-start helical strands in ice blue, gold, and coral. Each 3-start strand is a heteropolymeric thread consisting of alternating ArlB1 and 2. Protein atoms are displayed as spheres. Sugar moieties are shown as stick models.

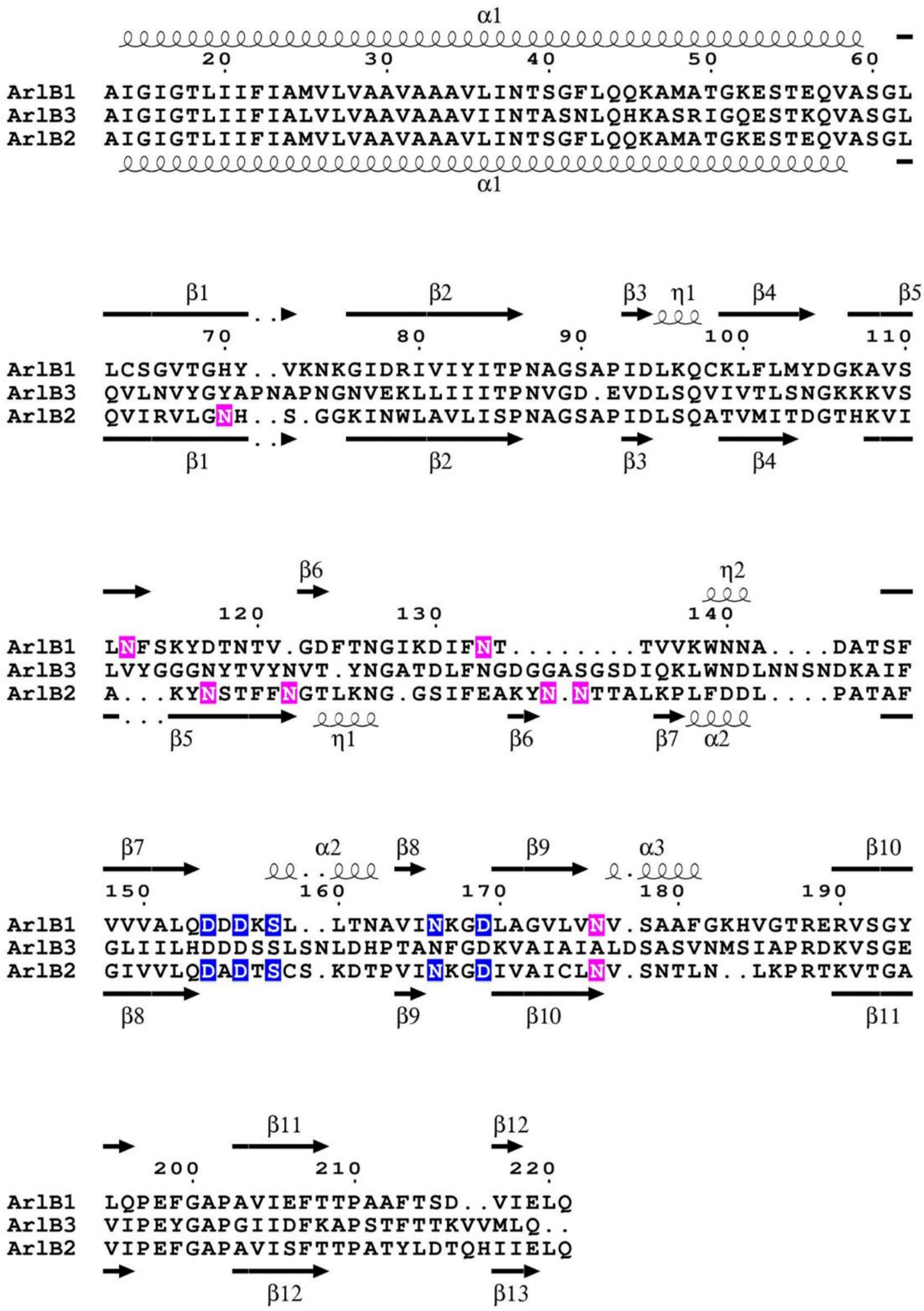

**Suppl. fig. 7|** Sequence alignment of *M. villosus* ArlB1, 2, 3 and secondary structure assignment of ArlB1 and ArlB2 archaellins. Secondary structure is indicated above and below the sequences as springs ("α" for α-helices and "η" for $3_{10}$-helices) or arrows ("β" for β-strands). N-glycosylated residues are highlighted in pink and metal ion binding residues are highlighted in blue.

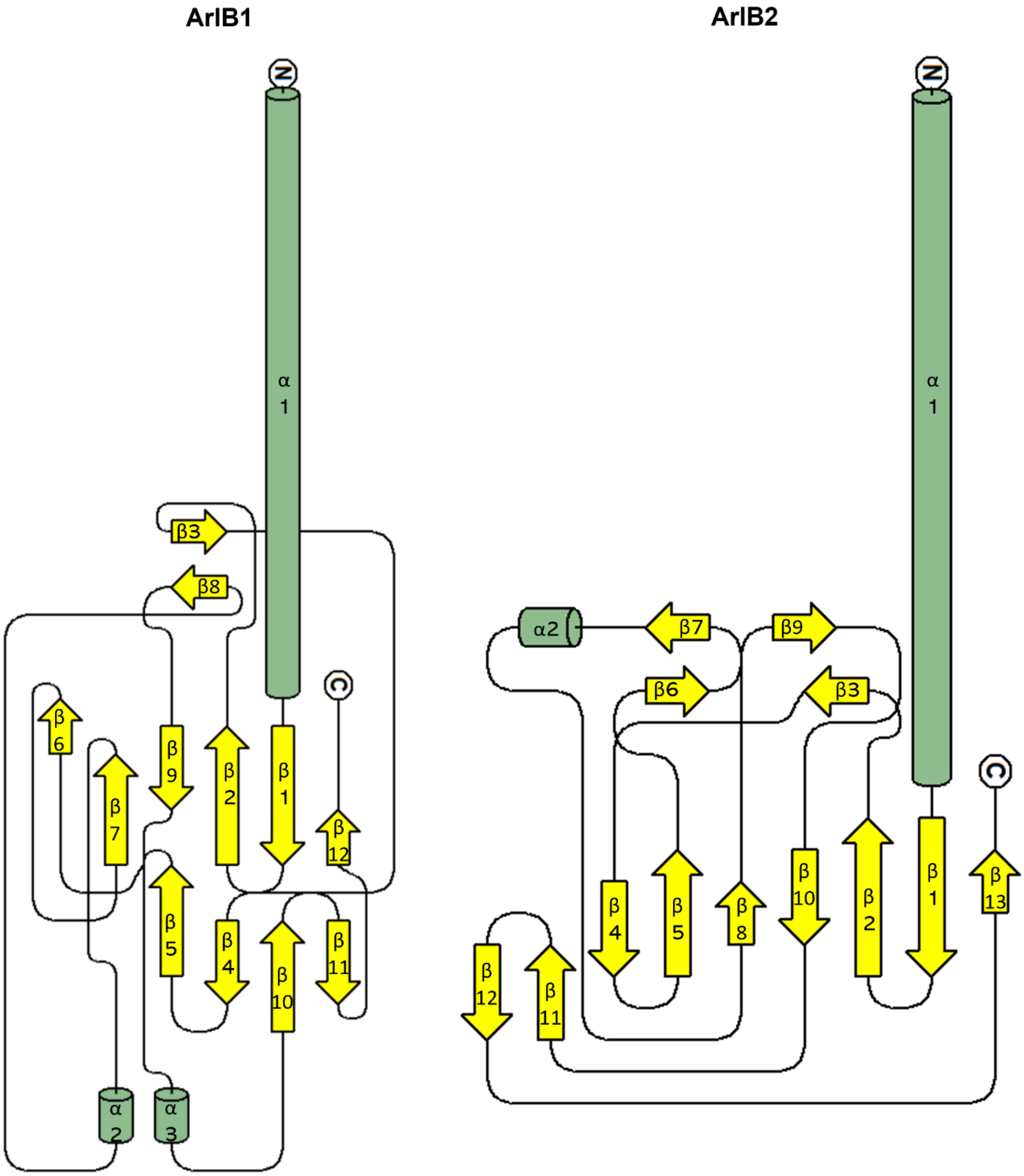


**Suppl. fig. 8|** Topology diagrams of ArlB1 and ArlB2.

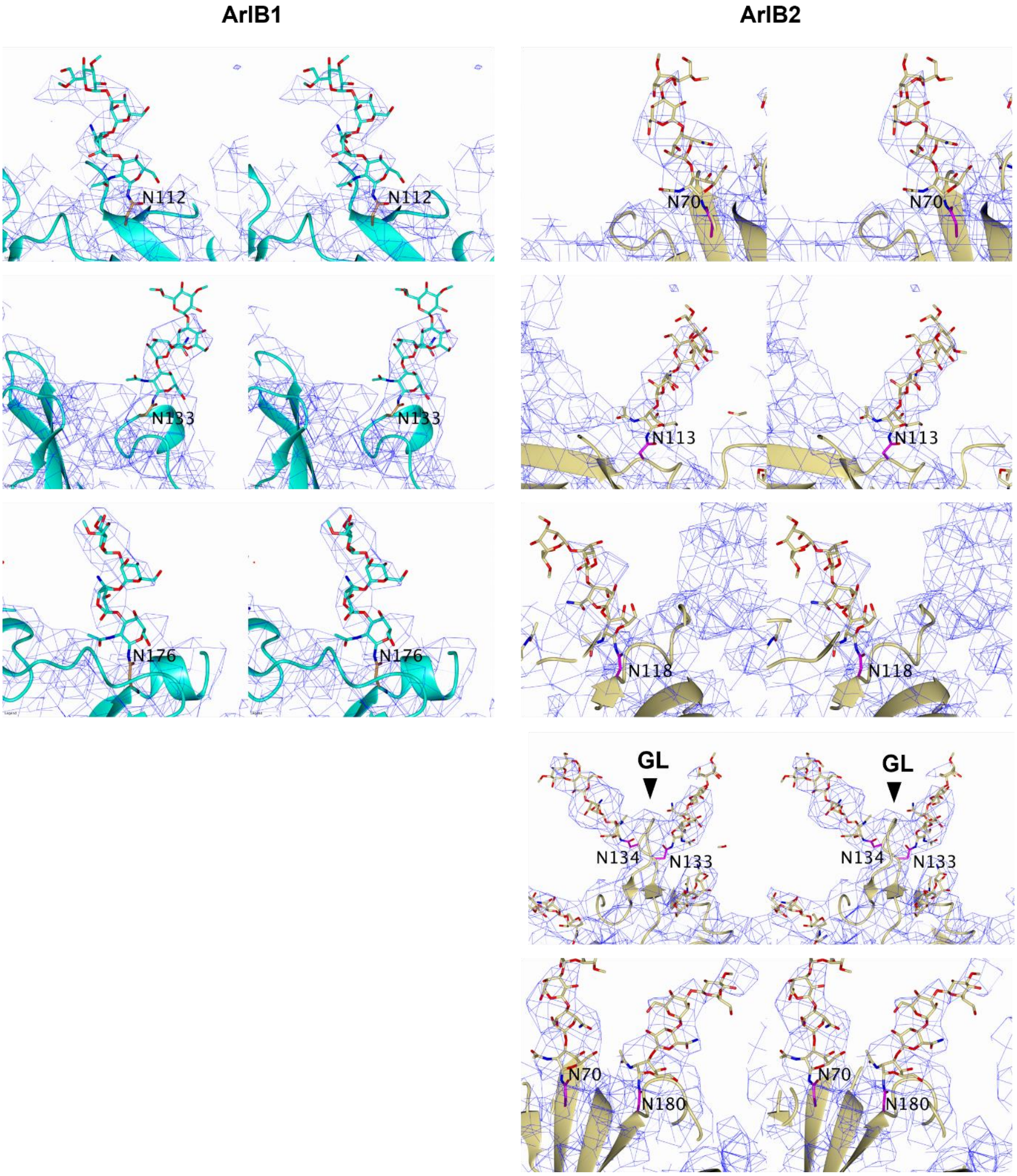


**Suppl. fig. 9|** Stereo diagrams showing the electron density for carbohydrate moieties of ArlB1 (protein ribbon and glycan carbons in cyan) and of ArlB2 (protein ribbon and glycan carbons in sand). Carbohydrates and side chains of glycosylated Asn residues are shown as stick models. Asn residues are additionally highlighted by differing carbon colours. GL, glycosylation loop.

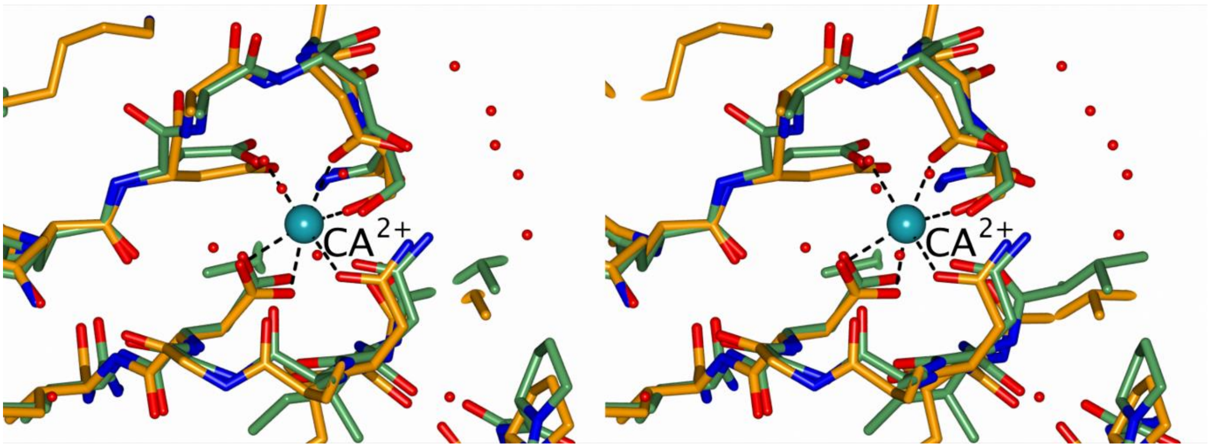


**Suppl. fig. 10|** Stereogram of the superposition of metal sites between *M. villosus* ArlB2 (orange) and *M. jannaschii* ArlB1 (green) (X-ray structure, PDB ID: 5ya6) shows a high degree of structural conservation.

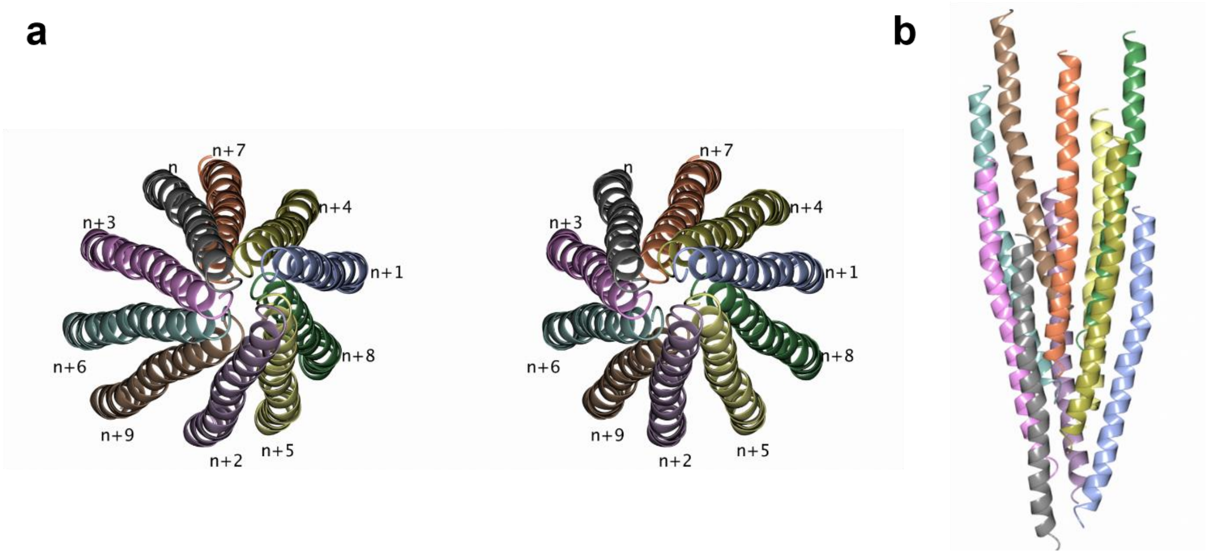


**Suppl. fig. 11|** Interactions between the tail helices in the archaellum filament. **a**, a stereo diagram showing ten tail domain helices viewed along the filament axis with each helix coloured differently and numbered in relation to the first one (n). **b**, top view of the helices in (a).

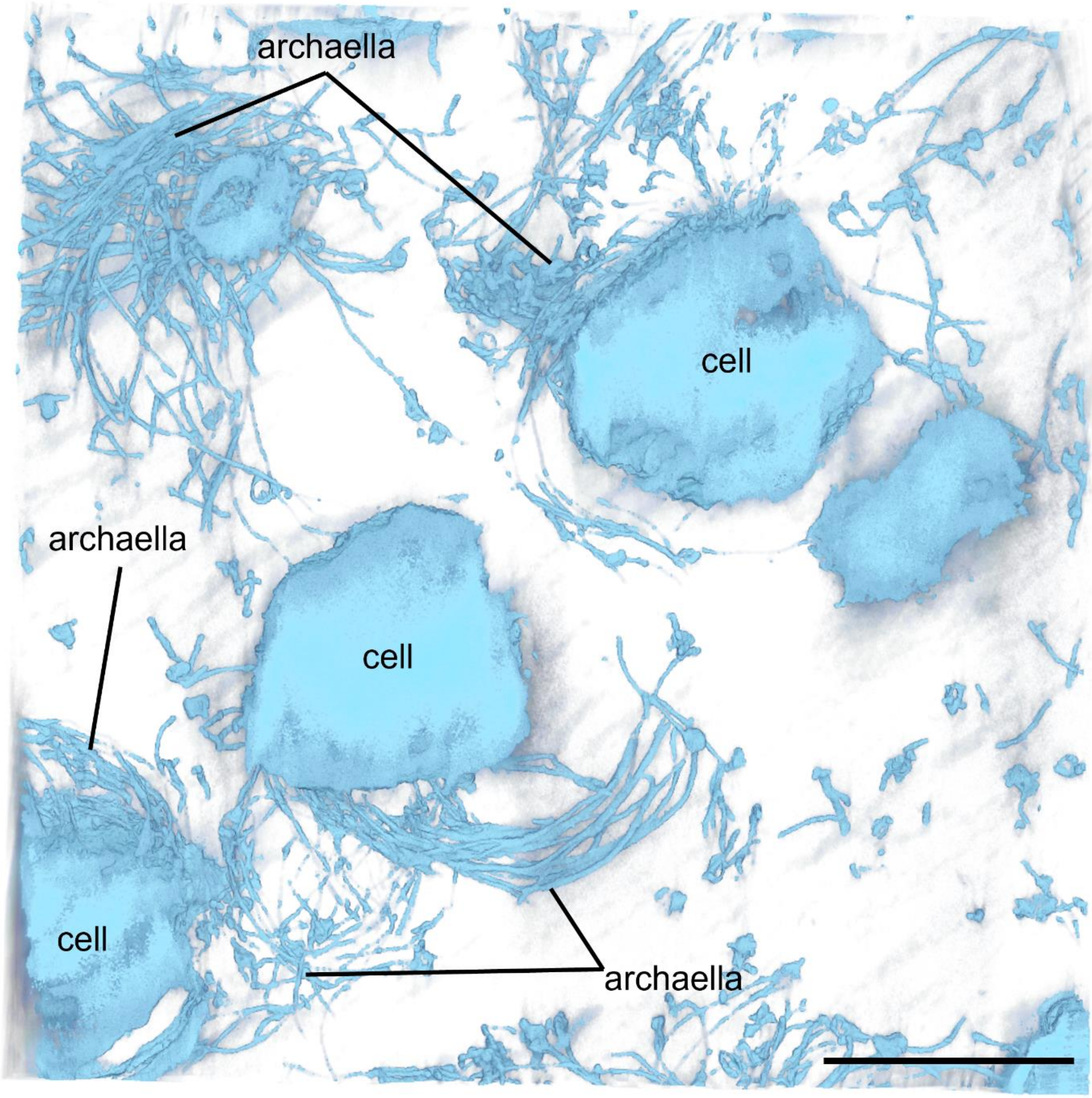


**Suppl. fig. 12|** STEM tomogram of freeze-substituted *M. villosus* cells. 3D representation was generated using solid representation in UCSF Chimera-X. Scale bar, 1 μm.

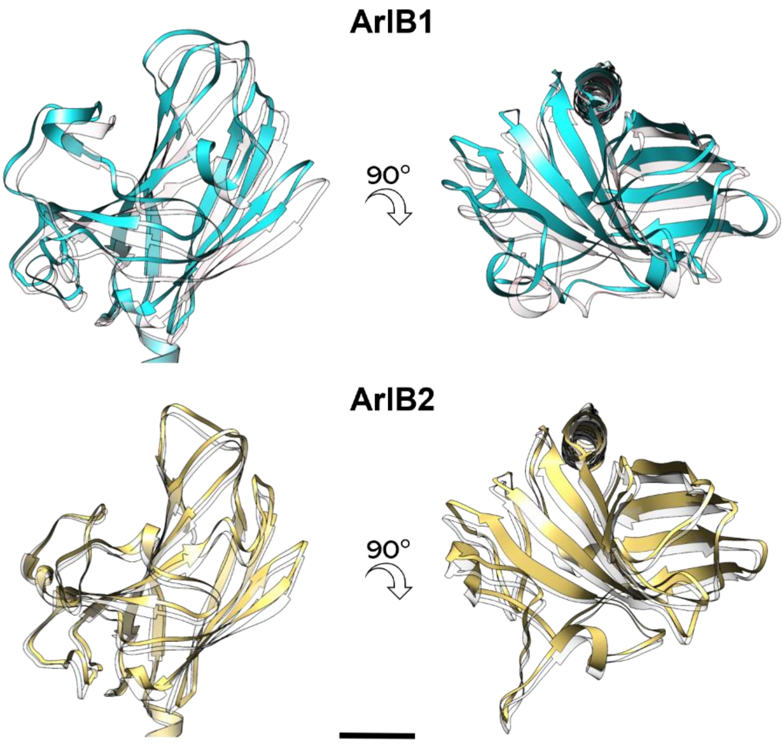


**Suppl. fig. 13|** Atomic models of one ArlB1 and one ArlB2 head domains showing their displacement during filament motion. The tail domains were aligned using *Coot*. The solid cyan/sand and transparent white models were fitted into the frame0 and frame19 maps of the cryoSPARC 3DVA respectively. The models highlight a displacement diagonal to the filament axis. Scale bar, 10 Å.

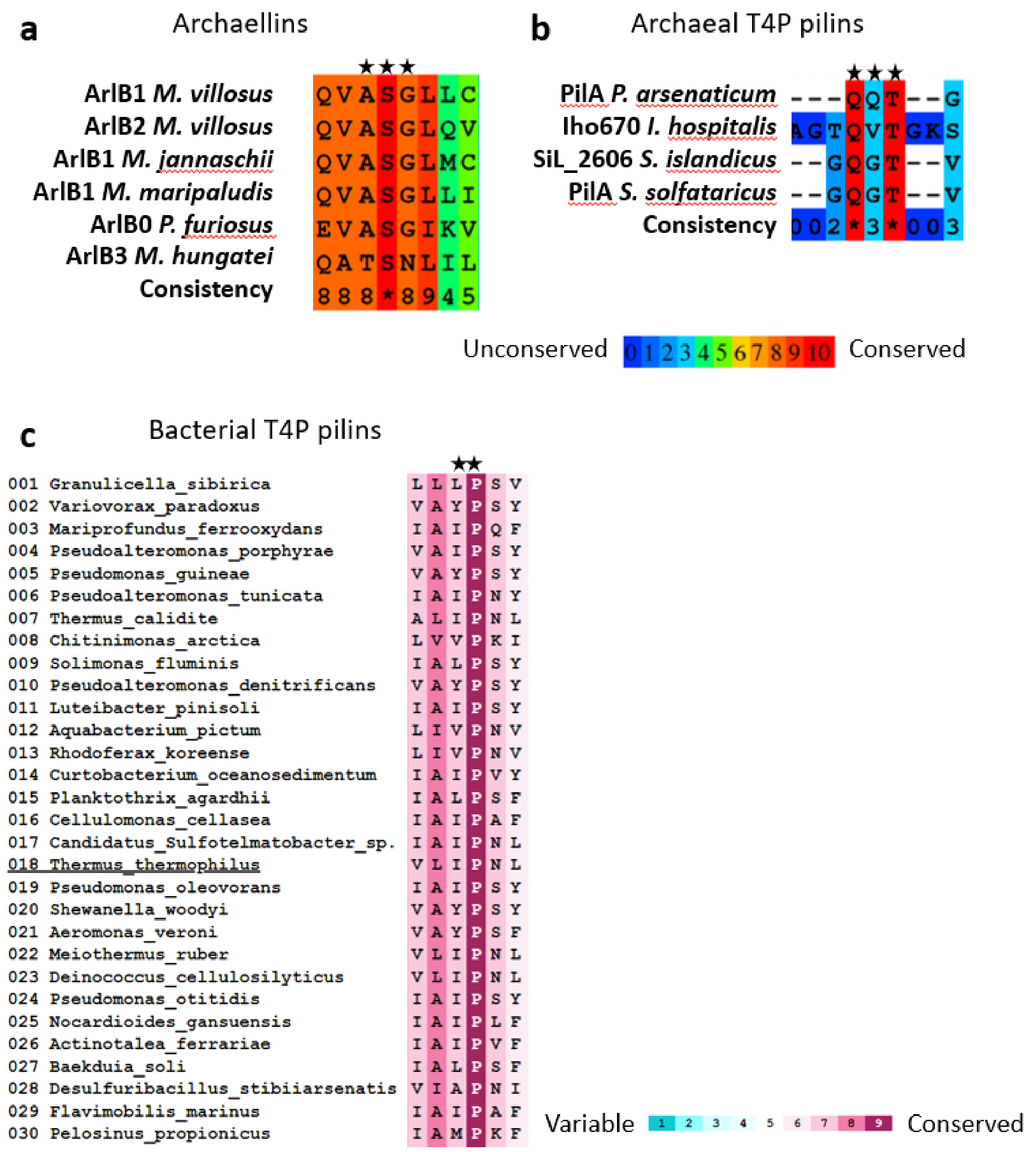


**Suppl. fig. 14|** Multi sequence alignment of the hinge region (starred) of archaellins (a), archaeal T4P pilins (b), and bacteria T4P pilins (c). For archaellins and archaeal pilins the MSA was performed in Praline and only species with known atomic structure were selected. For the bacterial T4P pilins the ConSurf server was used with the UNIREF90 database and *Thermus thermophilus* PilA4 (PDB ID: 6XXD) as template. The starred amino acids in (b) highlight the *T. thermophilus* PilA4 hinge region. All hinge regions are conserved in archaella and T4P, but their amino acids composition differs.

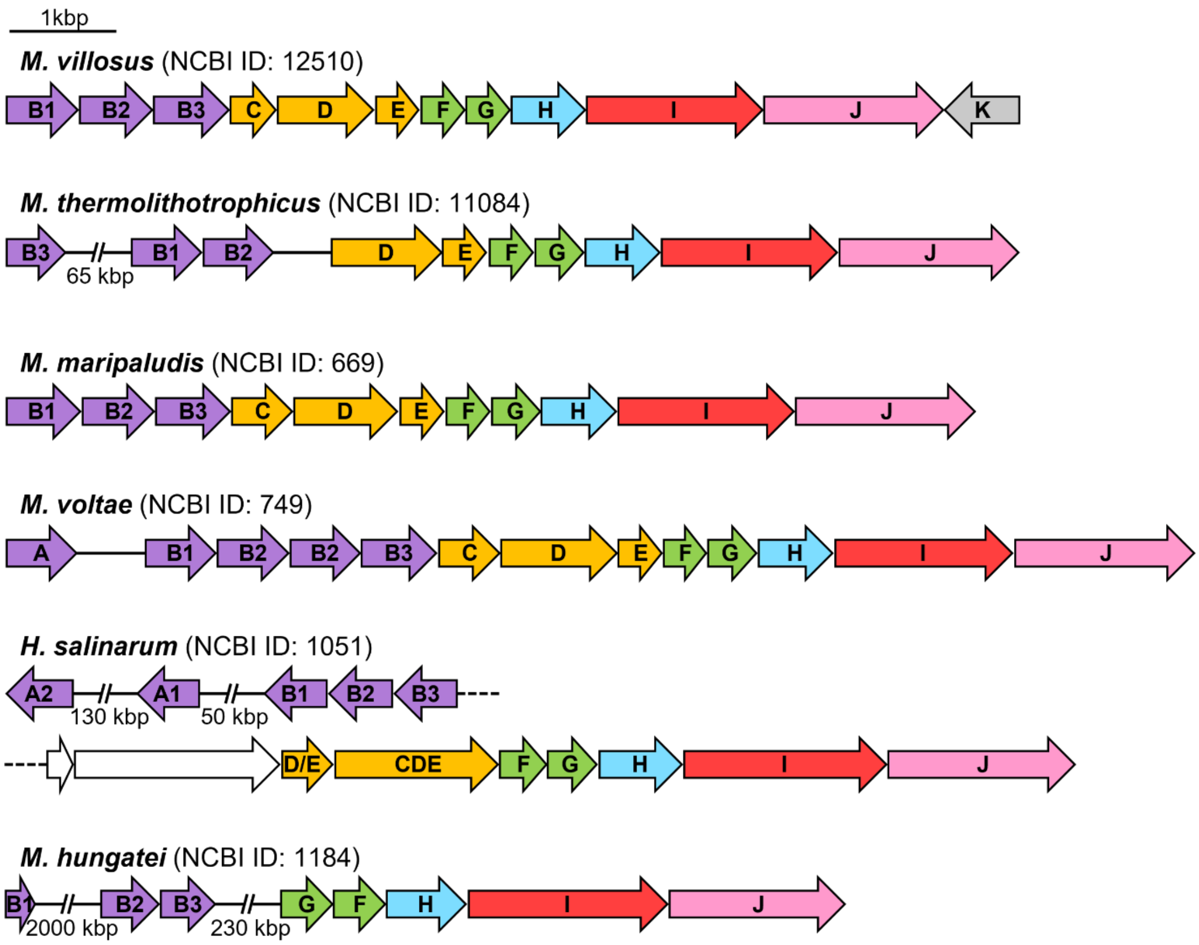


**Suppl. fig. 15|** Archaellum operons of archaea in which a hook at the base of the archaellum filament has been suggested. Purple: archaellins ArlA and B; orange: ring-forming proteins ArlC, D and E; green: periplasmic stator proteins ArlF and G; light blue: regulator protein ArlH; red: ATPase; pink: platform protein ArlJ; grey: prepilin peptidase ArlK; white: hypothetical protein.

**a**

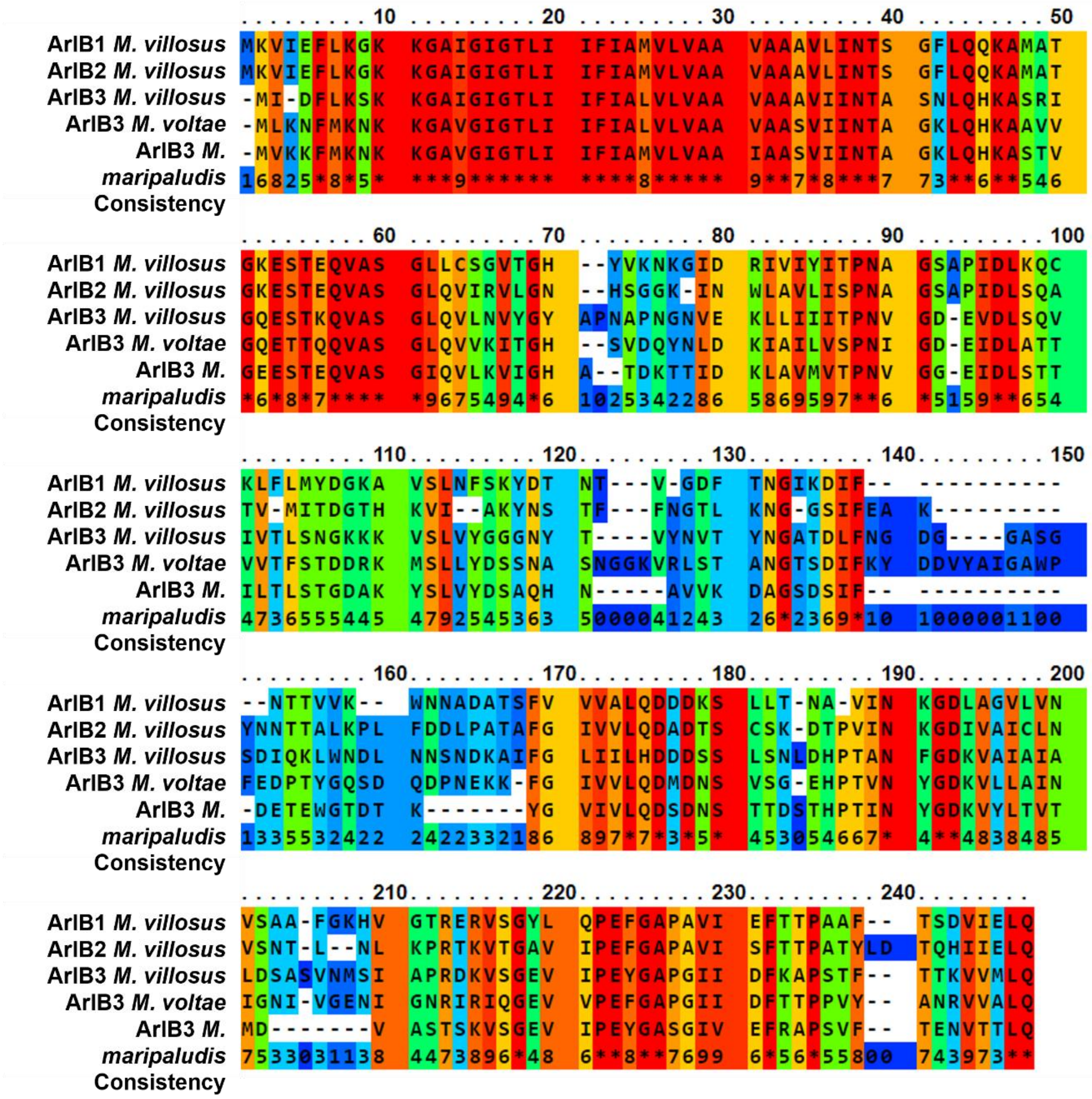


**b**

| | *M. villosus* ArlB1 | *M. villosus* ArlB2 | *M. villosus* ArlB3 | |
|---|---|---|---|---|
| *M. voltae* ArlB3 | 97 | 97 | 98 | sequence coverage (%) |
| | 41.5 | 45.6 | 48.3 | identity (%) |
| | 68.6 | 69.7 | 78.5 | similarity (%) |
| *M. maripaludis* ArlB3 | 99 | 99 | 98 | sequence coverage (%) |
| | 43.9 | 44.7 | 50.4 | identity (%) |
| | 70.6 | 71.9 | 75.4 | similarity (%) |

**Suppl. fig. 16| a**, sequence alignment of *M. villosus* ArlB1, 2, 3, *M. voltae* ArlB3 and *M. maripaludis* ArlB3 performed with Praline. **b**, sequence comparison between *M. villosus* ArlB1, 2, 3 with *M. voltae* ArlB3 and *M. maripaludis* ArlB3 using Blastp.

| | |
|---|---|
| **Data collection** | |
| Electron microscope | Titan Krios |
| Electron detector | Falcon III |
| Voltage (kV) | 300 |
| Defocus range (µm) | -2.3 to -1.1 in 0.3 increments |
| Pixel size ($Å^2$) | 1.39 |
| Total electron dose ($e^-/Å^2$) | 37 |
| Exposure time (s) | 1 |
| Number of fractions | 39 |
| Total movies | 2,759 |
| **3D reconstruction** | |
| Final particles | 399,178 helical segments |
| Resolution (masked FSC=0.143), Å | 3.08 |
| B factor | -137.584 |
| EMDB accession # | 12875 |
| **Model Refinement** | |
| PDB ID | 7OFQ |
| Model resolution (FSC = 0.50/0.143), Å | 3.35 / 3.06 |
| Model refinement resolution, Å | 3.08 |
| Non-hydrogen atoms (overall/protein/metal/glycan) | 80,691/70,194/45/10,452 |
| Number of monomers (overall/ArlB1/ArlB2) | 45/23/22; 1.5 full helical turns |
| **RMS deviations** | |
| Bond length (Å) | 0.010 |
| Bond angle (°) | 2.33 |
| **Ramachandran plot** | |
| Favoured (%) | 95.00 |
| Allowed (%) | 4.99 |
| Outliers (%) | 0.01 |
| **Validation** | |
| Rotamer outliers (%) | 1.96 |
| Molprobity score | 1.72 |
| Clash score | 3.65 |

**Suppl. table 1|**Statistics of data collection, 3D reconstruction and validation.

| Parameter/ Contact type | BSA[a] $Å^2$ | Hydrogen bond | Salt bridge | dG_diss kcal/mol |
|---|---|---|---|---|
| n + 3 ArlB1 - 2 | 3,356 | 13 | 3 | 20.4 |
| n + 3 ArlB2 – 1 | 3,414 | 8 | 2 | 16.2 |
| n + 3 ArlB1 - 2 (head domains only) | 1,529 | 12 | 2 | 0.9 |
| n + 3 ArlB2 - 1 (head domains only) | 1,686 | 7 | 1 | - |
| n+7 ArlB1 - 1 | 1,794 | 0 | 4 | 1.6 |
| n+7 ArlB1 - 2 | 1,663 | 0 | 2 | 1.7 |
| n+7 ArlB2 - 1 | 1,795 | 0 | 5 | 4.7 |
| n+7 ArlB2 - 2 | 1,731 | 0 | 5 | 3.7 |

[a] BSA - buried solvent accessible surface area

**Suppl. table 2|** Protein contacts in the heteropolymeric archaellum of *M. villosus*.

**Suppl. movie 1|** CryoEM map of the *M. villosus* archaellum filament showing alternating subunits.

**Suppl. movie 2|** CryoEM map and atomic model of the *M. villosus* archaellum filament.

**Suppl. movie 3|** Morphing of the 20 cryoEM maps of the cryoSPARC 3D variability analysis showing the *M. villosus* archaellum flexibility.

**Suppl. movie 4|** Atomic model showing the flexibility of the *M. villosus* archaellum filament.

**Suppl. movie 5|** Atomic model of two opposite n+10 sets of protein monomers showing the flexibility of the tail domains.

**Suppl. movie 6|** Atomic model in backbone representation showing the flexibility of the head domain of ArlB1 along a full filament turn of a homopolymeric pseudo-strand.

**Suppl. movie 7|** Atomic model in backbone representation showing the flexibility of the head domain of ArlB2 along a full filament turn of a homopolymeric pseudo-strand.
